# scDILT: a model-based and constrained deep learning framework for single-cell Data Integration, Label Transferring, and clustering

**DOI:** 10.1101/2023.10.09.561605

**Authors:** Xiang Lin, Jianlan Ren, Le Gao, Zhi Wei, Junwen Wang

## Abstract

The scRNA-seq technology enables high-resolution profiling and analysis of individual cells. The increasing availability of datasets and advancements in technology have prompted researchers to integrate existing annotated datasets with newly sequenced datasets for a more comprehensive analysis. It is important to ensure that the integration of new datasets does not alter the cell clusters defined in the old/reference datasets. Although several methods have been developed for scRNA-seq data integration, there is currently a lack of tools that can simultaneously achieve the aforementioned objectives. Therefore, in this study, we have introduced a novel tool called scDILT, which leverages a conditional autoencoder and deep embedding clustering to effectively remove batch effects among different datasets. Moreover, scDILT utilizes homogeneous constraints to preserve the cell-type/clustering patterns observed in the reference datasets, while employing heterogeneous constraints to map cells in the new datasets to the annotated cell clusters in the reference datasets. We have conducted extensive experiments to demonstrate that scDILT outperforms other methods in terms of data integration, as confirmed by evaluations on simulated and real datasets. Furthermore, we have shown that scDILT can be successfully applied to integrate multi-omics single-cell datasets. Based on these findings, we conclude that scDILT holds great promise as a tool for integrating single-cell datasets derived from different batches, experiments, times, or interventions.

## INTRODUCTION

The utilization of single-cell RNA sequencing (scRNA-seq) technology allows for the comprehensive profiling of various biological activities at the cellular level, including gene expression, protein expression, and chromatin accessibility. With the significant proliferation of available datasets, researchers are increasingly interested in conducting integrated analyses of datasets originating from diverse batches, tissues, and samples. However, the presence of batch effects poses a big challenge as it hinders the comparability and compatibility of different datasets[1]. Hicks et al. indicated that batch effects in scRNA-seq experiments occur when cells are cultured, captured, and sequenced separately[2] and two totally identical experimental designs are generally impossible to be achieved[3] [4]. Thus, before analyzing different batches of datasets, it is of great importance to correct the batch effect induced by these technical factors[2].

Many methods have been developed for batch effect removal in scRNA-seq data[5] [6] [7] [8] [9] [10]. Most of them correct the batch effect in a low-dimensional representation of original data. The integrated latent space can be used for many downstream analyses, such as clustering analyses and trajectory analyses. Seurat 3.0[9] corrects batch effect by finding anchor pairs between two batches of data. It relies on the mutual nearest neighbor (MNN) approach[11] which can only integrate two batches of data at a time. The performance is influenced by the order in which the input data is processed, and Seurat becomes computationally expensive in terms of both time and space when dealing with a large number of cells. Furthermore, when combining a pre-clustered and annotated dataset with one or more new datasets, Seurat operates in an unsupervised manner, leading to potential perturbations in the pre-defined cell types. This outcome is less desirable for biologists, as the annotation process is labor-intensive and exhaustive. Biologists aim to maintain the integrity of the original clusters after integration and incorporate new cells into the existing clusters without disrupting the established annotations.

Since the annotation process is very exhaustive, many studies have been designed to transfer labels from a reference dataset to the query datasets. Some models employ supervised approach to predict the cell type of the query datasets[12]. However, when the reference and query datasets are under different conditions (such as pre- and post-treatment), supervised methods can only assign new cells to the existent cell types and cannot discover new cell types in the query datasets. On the other hand, unsupervised clustering methods, such as scGCN[13], CarDEC[14] and DESC[7], are also unsatisfactory because the prior information (annotated cell types in the reference dataset) cannot be used to guide the analyses of the integrated data, such as the representation learning and clustering process. Seurat V3 also works in an unsupervised manner. However, the labels are directly transferred between the anchors in reference and query datasets.

To tackle the aforementioned challenges, we have developed a semi-supervised model known as Single-Cell Deep Data Integration and Label Transferring (scDILT). This model aims to integrate diverse batches of data while simultaneously transferring labels from a reference dataset to one or more query datasets. The scDILT framework leverages a conditional autoencoder (CAE) to accomplish this task. The CAE receives the concatenated count matrix of multiple datasets, along with a vector indicating the batch IDs. By doing so, the CAE learns a latent representation of the integrated datasets while effectively mitigating the impact of batch effects on the data. Additionally, we have incorporated deep embedding clustering (DEC) to the integrated latent space generated by CAE, aiming to further enhance the removal of batch effects. To use the true label of reference data to guide the latent representation learning and clustering of query data, we build cell-to-cell constraints and implement them on the latent space[15].

In the clustering process, scDILT incorporates two types of constraints to ensure effective integration. Firstly, it utilizes homogenous constraints, which preserve the pre-defined clustering or cell-typing patterns observed in the reference datasets. This ensures that the integrated results maintain the original cluster structure and cell type annotations from the reference datasets. Secondly, scDILT employs heterogeneous constraints to accurately map the cells from the new datasets to the pre-defined cell types or clusters in the reference datasets. By leveraging these constraints, scDILT ensures that the integration process accurately assigns the cells from the new datasets to the appropriate cell types or clusters based on the information provided by the reference datasets or datasets that are previously annotated.

Upon completion, scDILT generates an integrated latent space representing the input datasets along with predicted labels for all cells. Through extensive experimentation, we have demonstrated that scDILT surpasses existing methods in terms of data integration, label transferring, and the discovery of new cell types. Furthermore, we have successfully applied scDILT to integrate scRNA-seq and scATAC-seq data in two Single-cell Multiome ATAC and Gene Expression datasets. Based on the clustering results obtained from scDILT, we conducted differential expression analyses. The outcomes of these analyses illustrate the superior performance of scDILT in integrating multi-omics data. Consequently, we conclude that scDILT holds substantial promise as a tool for jointly analyzing multiple scRNA-seq datasets.

## METHODS AND MATERIALS

### Data preprocessing

Our model can be used to integrate and cluster scRNA-seq datasets from different batches or experiments. For example, we have two datasets 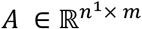and 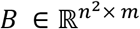from different experiments, and we want to integrate them. We first select the top 2000 highly variable genes (HVGs) for the concatenated datasets of A and B 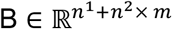. In this way, the filtered data A and B will have the same genes. We then separate the raw count data of A and B and preprocess and normalize them by the Python package SCANPY[16]. Specifically, the genes with no count are filtered out. The counts of a cell are normalized by a size factor *s*_*i*_, which is calculated as dividing the library size of that cell by the median of the library size of all cells. In this way, all cells will have the same library size and become comparable. Then, the counts are logarithm transformed and scaled to have unit variances and zero means. The treated count data are used in our denoising conditional autoencoder model. The raw count matrices are used to calculate the ZINB loss[17] [18]. Since the selected HVGs are the same for A and B, we finally concatenate the normalized data matrix of A and B as the input for our model. This approach can also be used to preprocess more than two datasets.

### Conditional denoising autoencoder

The denoising autoencoder is a neural network for learning a nonlinear representation of data [19]. It receives corrupted data with artificial noises and reconstructs the original data with an encoder and a decoder[20]. It is generally used to learn a robust latent representation for noisy data. Here we employ the denoised autoencoder to the scRNA-seq data which is highly noised. Let’s denote the preprocessed counts data as *X* and the corrupted data as *X*_*c*_, formally:

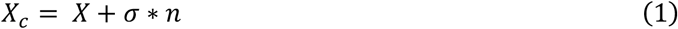

where *n* is the artificial noise in standard Gaussian distribution (with mean=0 and variance=1), and *σ* controls the weights of *n*. We set *σ* as 2.5.

Next, we use an autoencoder to reduce the dimension of count data. Encoders (*E*) and decoders (*D*) are both multi-layered fully connected neural networks. Denoting the latent space as Z and the reconstructed count matrix as *X*^′^, the encoder is Z = *E*_w_(*X*_*c*_) and the decoder is *X*^(^ = *D*_w(_(Z), where *w* and *w*^′^ are the learnable weights for encoder and decoder, respectively. Conditional autoencoder (CAE) and conditional variational autoencoder have been used to integrate the data from different batches[21]. Based on the traditional autoencoder model, we add a matrix B on the input of encoder and decoder. B is the one-hot coding from a batch vector b of each cell. If there are M batches in b, the dimension of B would be *N* × *M*. So, the encoder becomes Z = *E*_w_(*X*_*c*_ ⊙ *B*) and the decoder becomes *X* (= *D*_w(_(Z ⊙ *B*) where ⊙ means the concatenation of two matrices. Note, *X*_*c*_ and Z are only used for model training. We perform the downstream analyses based on the latent space Z* which is obtained from the well-trained model without adding artificial noise. The ELu activation function[22] is used for all the hidden layers in the encoder and decoder except the bottleneck layer.

### ZINB loss

We employ a zero-inflated negative binomial (ZINB) model in the reconstruction loss function to characterize the zero-inflated and over-dispersed count data[23]. This distribution is parameterized by three coefficients, namely the mean (*μ*_*ij*_), the dispersion (*θ*_*ij*_), and the probability of dropout events (zero mass, π_*ij*_). The raw count data, not the normalized data, is used in the ZINB model[17] [18] [23]. Let *X*_*ij*_ be the count for cell i and gene j in the raw count matrix. The NB distribution is defined as :

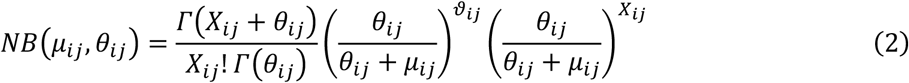

Then, with an additional coefficient π_*ij*_, ZINB is defined as :

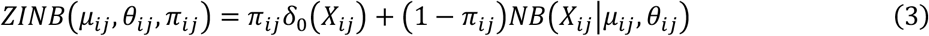

The ZINB-based reconstruction loss of the autoencoder is defined as:

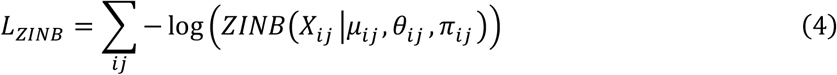

We add three independent fully connected layers *M, Θ*, and *Π* as the parameter layers after the last hidden layer of the decoder which outputs the reconstructed matrix *X*^′^. The parameter layers are defined as:

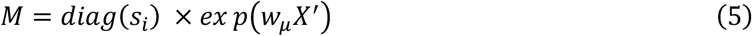

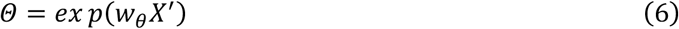

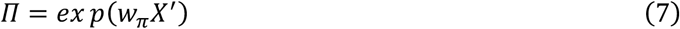

where *M, Θ*, and *Π* are the matrix of estimated mean, dispersion, and drop-out probability for the ZINB loss. *w*_*μ*_, *w*_*θ*_, and *w*_*π*_ are the learnable weights for them, respectively. The size factor *s*_*i*_ for the cell i was calculated in the preprocessing step.

The sizes of layers are set to (256, 128, 64, 32) for the encoder and (32, 64, 128, 256) for the decoder. The overall architecture of the scDILT model is shown in **Fig. 1**.

**Figure 1.**
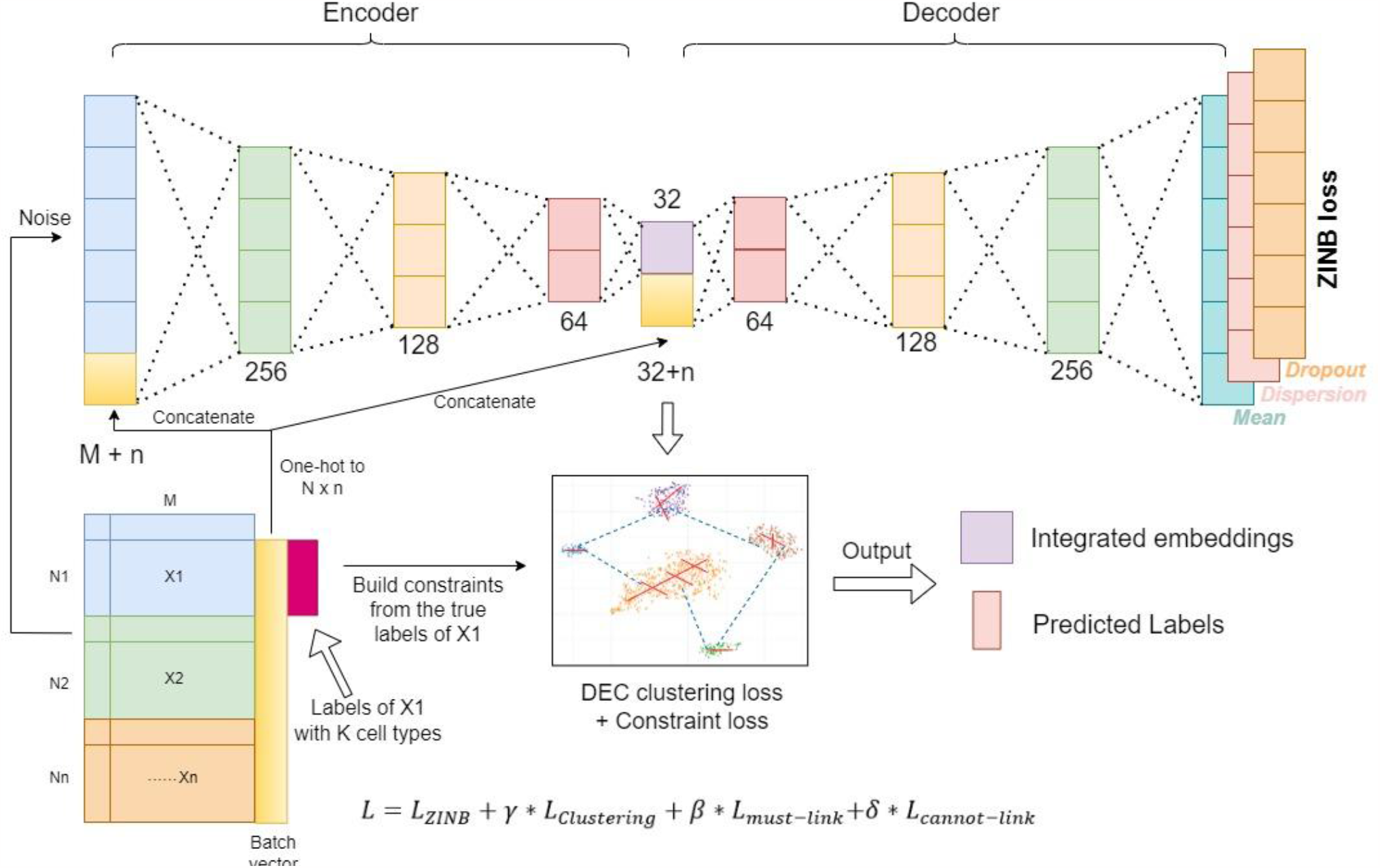
The model architecture of scDILT. This model employs a conditional autoencoder structure to integrate the data from different batches. A batch vector is one-hot encoded and concatenated to the input of encoder and decoder. If one or more datasets are used as the reference (with annotated cell types), the cell-to-cell constraints will be built based on the labels of these data and implemented on the bottle-neck layer Z of the autoencoder. Then the deep embedding clustering will be performed on Z. scDILT will output the integrated embeddings of the multiple batches of data and the predicted labels of the cells in all the batches.

### Deep embedded clustering

Our model has two learning stages, a pretraining stage and a clustering stage. In the pretraining stage, we train the autoencoder without considering the clustering loss and the constraint loss (see details below). In the clustering stage, we simultaneously optimize the autoencoder and the clustering results under the guidance of constraints. We perform unsupervised clustering on the latent space of the autoencoder[24]. Our autoencoder transfers the input matrix to a low dimensional space *Z*. The clustering loss is defined as the Kullback-Leibler (KL) divergence between the soft label distribution *Q’* and the derived target distribution *P’*.

Formally, Q’ is defined as:

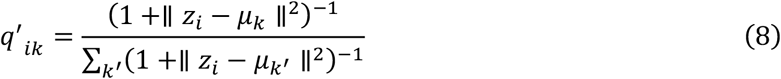

Where _4_^′^_*i*4_ measures the similarity between _4*i*_ and cluster center *μ*_4_ by Student’s t-kernel[25] and the cluster center *μ*_4_ is initialized by applying a K-means on the latent space Z from the pretraining stage, and then updated per batch in the clustering stage. The target distribution _P_^′^ which emphasizes the more certain assignments is derived from *Q’*. Formally _P_^′^_*ik*_ϵ*P*^′^ is defined as:

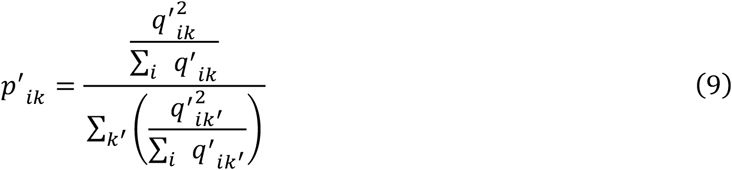

During the training process, _P_^′^ and clustering loss are calculated per batch and _P_^′^ is updated per epoch. Then, the clustering loss is calculated as:

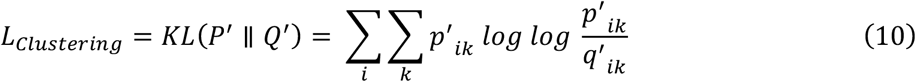

### Autoencoder with pairwise constraints Homogeneous constraints

When one or more of the input datasets have annotated label/cell types, we use the information of these labels to guide the clustering of the unlabeled cells. Based on the autoencoder architecture, we add pairwise constraints of cells[15] on the latent space according to the labels of cells in the reference datasets. We employ the must-link (ML) constraints to push two cells to have similar soft labels if they are in the same cell types. On the other hand, we employ the cannot-link (CL) constraints to pull two cells to have distinct soft labels if they are in the different cell types. The constraints loss of must-link is defined as:

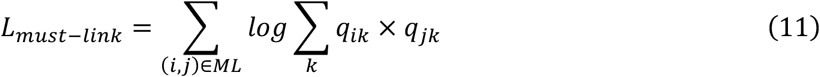

where *a*_*ik*_ is the soft label of cell i for cluster k as described in the clustering section above. The constraints loss of cannot-link is defined as:

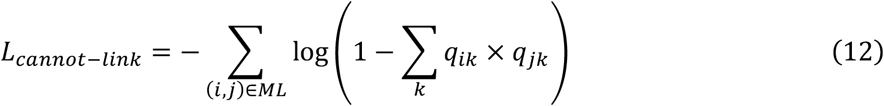

The number of constraints can be set according to the cell numbers. In this study, we set 3000 must-link constraints and 9000 cannot-link constraints. By adding the constraints in the autoencoder model, the clustering results of the reference dataset(s) can be retained in the new clustering analysis and be transferred it to the new datasets. Besides, although the constraints are only built between the cells in the reference dataset(s), they can help to improve the clustering performance of the unlabeled datasets by locating the centroids of each cluster.

### Heterogenous constraints

To connect the cells in different datasets, we employ the algorithm in Seurat V3 to calculate the anchor cells in two different datasets. Briefly, Seurat uses canonical correlation vectors (CCV) to project the two datasets into a correlated low-dimensional space. Then mutual nearest neighbors in the two datasets are identify based on the L2-normalized CCV. We implement these algorithms by using Seurat (V4.1.0). Firstly, the raw count data of each dataset are normalized, and the top 2000 highly variable genes are selection for each. Then, the functions SelectIntegrationFeatures() is used to find the anchors between the two datasets. These anchors will be used as the must-links in scDILT(Table S5-S10). Combining the pairwise constraint loss, reconstruction loss, and clustering loss, the total loss of the scDILT is:

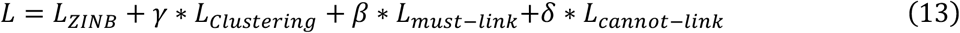

Where 𝒳, *β* and *δ* are the coefficients for the clustering loss, must-link loss (includes the loss from homogeneous and heterogeneous constraints), and cannot-link loss respectively. In the experiments of this study, we set 𝒳, *β* and *δ* as 1, 0.5 and 1.5.

### Model implementation

This model is implemented in Python3 using PyTorch (Paszke, et al., 2017). Adam with AMSGrad variant (Kingma and Ba, 2014; Reddi, et al., 2018) with an initial learning rate = 0.001 is used for the pretraining stage. Adadelta (Zeiler, 2012) with learning rate 1.0 is used for the clustering stage. The batch size is set as 256. We pretrain the autoencoders for 400 epochs before entering the clustering stage. In the beginning of the clustering stage, we initialize *K* centroids by the k-means algorithm. During the clustering stage, all loss functions (*L*_*ZINB*_, *L*_*Clustering*_, *L*_*must-link*_ *L*_*cannot-link*_) are optimized simultaneously, and the centroids are also continuously updated by the learning process. The soft label distribution *Q’* is calculated in each batch and the derived target distribution *P’* is updated after each epoch. The convergence threshold for the clustering stage is that less than 0.1% of labels are changed per epoch. All experiments of scDILT in this study are conducted on NVIDIA GeForce RTX 3090._4_The links we used is listed in Table 2. In general, the number of total links, including homogenous link (ML1), heterogenous link (ML2) and cannot link (CL), is approximately same as cells numbers, and the ratio of ML1 plus ML2 to CL is around 1 to 3. We find this combination reaches the robust and high clustering performance in all datasets (Table 2). For other model tests, we discuss in the results part.

### Evaluation metrics for clustering performance

In a well-integrated data, the cells in the same cell type from different datasets can be assigned into the same cluster. Thus, we evaluate the performance of data integration by measuring the clustering performance. Adjusted Rand Index (ARI)[26] and Normalized Mutual Information (NMI)[27] are used as metrics to evaluate the clustering performance of different methods.

Adjusted Rand Index measures the agreements between two sets U and G. Assuming a is the number of pairs of two cells in the same group in both U and G; b is the number of pairs of two cells in different groups in both U and G; c is the number of pairs of two cells in the same group in U but in different groups in G; and d is the number of pairs of two cells in different groups in U, but in the same group in G. Formally, the ARI is defined as:

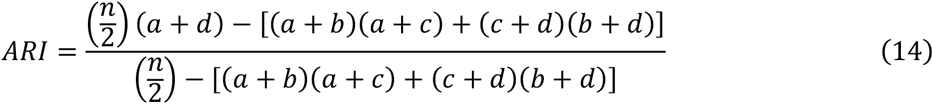

Let U = {U1, U2, …, U_tu_ } and G = {G1, G2, …, G_tg_} be the predicted and ground truth labels of a dataset. NMI is calculated based on the mutual information between U and G (I(U,G)) and their entropies. Formally, I(U,G) is defined as:

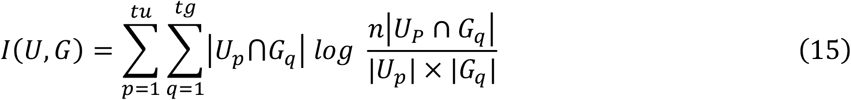

And the entropy of U (H(U)) and U (H(G)) are formalized as:

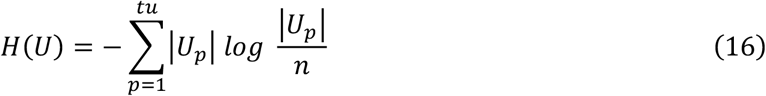

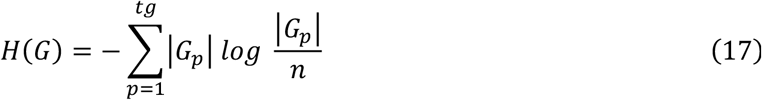

Then, NMI is defined as:

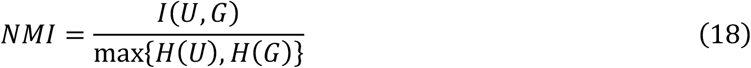

### Real datasets

The first set of data are downloaded from GEO with accession number GSE150599 (https://www.ncbi.nlm.nih.gov/geo/query/acc.cgi?acc=GSE150599). Also, we only use the scRNA-seq data from it. This dataset contains two spleen data from two mice. Each data has two batches. The cell types are provided by the author. The second set of data is from GSE131907 (https://www.ncbi.nlm.nih.gov/geo/query/acc.cgi?acc=GSE131907) and GSE123904 (https://www.ncbi.nlm.nih.gov/geo/query/acc.cgi?acc=GSE123904). Same as last, only scRNA-seq data is used from them. The cell types are provided from the author. The third set of data are from GEO with accession number GSE132802 (https://www.ncbi.nlm.nih.gov/geo/query/acc.cgi?acc=GSE132802). We use four scRNA-seq data from this dataset three tissues including: 1) PBMC from a healthy person (https://satijalab.org/signac/articles/pbmc_vignette.html); 2) PBMC from the patient with Drug-induced hypersensitivity syndrome/drug reaction with eosinophilia and systemic symptoms (DiHS/DRESS) but without treatment; 3) PBMC from the patient with sulfamethoxazole (SMX) and trimethoprim (TMP) treatment (BACT); 4) PBMC from the patient with SMX, TMP, and tofacitinib (TOFA) treatment. The scRNA-seq data from a normal person is provided by 10X genomics and annotated by Satija lab (https://www.dropbox.com/s/kqsy9tvsklbu7c4/allen_brain.rds?dl=0). The fourth set of data is the real Single-cell Multimode ATAC Gene Expression (SMAGE-seq) dataset from human PBMC downloaded from the 10X Genomics website (https://www.10xgenomics.com/resources/datasets). This dataset contains both the scRNA-seq data and the scATAC-seq data for the same cells. Since the peak matrix from scATAC-seq data is very sparse, we integrate the peaks into the gene regions by using Signac (1.4.0) in R. Details of each dataset are shown in **Table 1**.

**Table 1.**
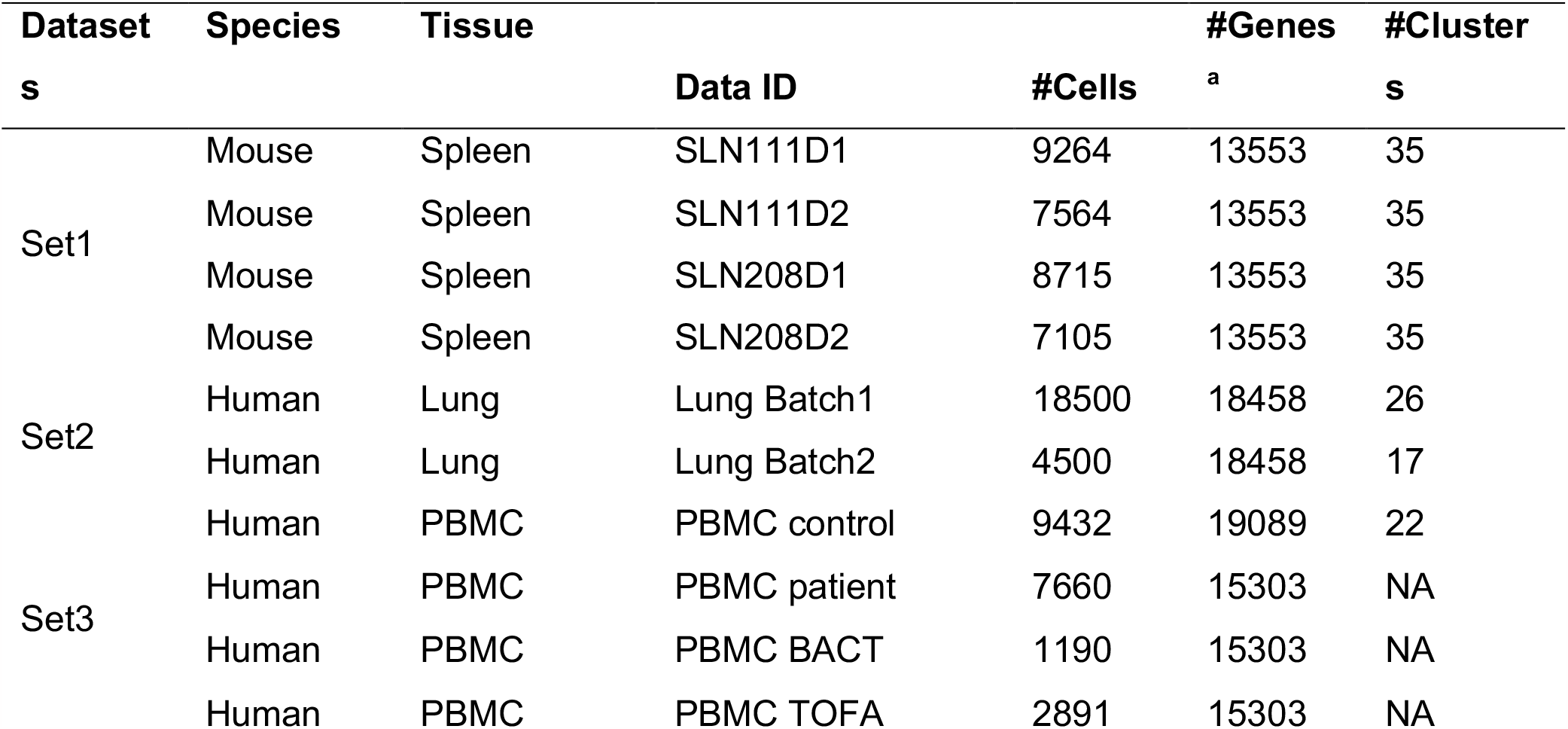
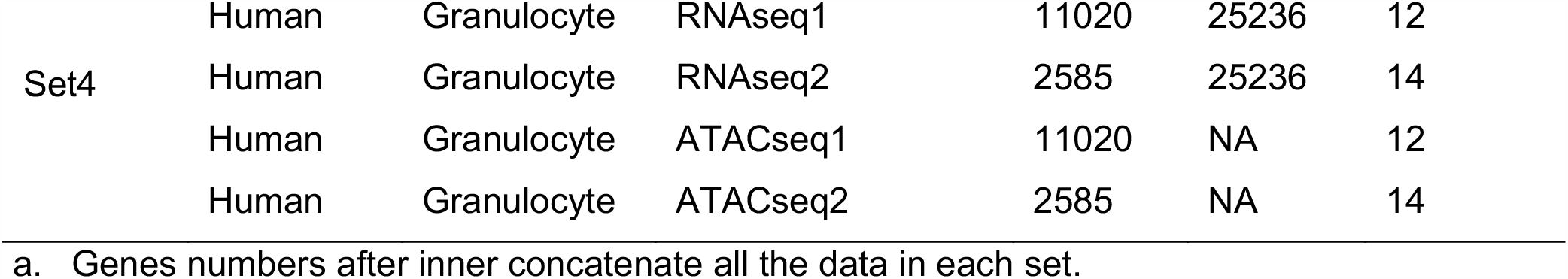
Summary of the real datasets.

**Table 2.** Link number used in experiments.

|  | ML1 | ML2 | CL |
| --- | --- | --- | --- |
| Simulation | 1000 | 500 | 4000 |
| SLN111 | 1500 | 1500 | 9000 |
| SLN208 | 1500 | 1500 | 9000 |
| Lung cancer | 2000 | 1000 | 9000 |
| PBMC 4 batches | 1000 | 1000 | 6000 |
| 10X rnaseq | 2000 | 2000 | 9000 |
| 10X atacseq | 2000 | 2000 | 9000 |

### Competing methods

We compared our method with three state-of-art methods: 1) Seurat v3 with batch effect removal (Stuart, et al., 2020). The resolution parameter for estimating the number of clusters (K) is tuned to get a similar K with the real K. 2) DESC (v2.1.1)[7]. 3) CarDEC[14]. All the methods employ the same data preprocessing/normalization and feature selection approaches. Top 2000 variable genes are selected before analysis. Other parameters are kept as default.

### Simulation

For the generation of simulated datasets, we utilized the R package Splatter (v1.18.2). The simulation parameters were estimated from a real dataset, GSE126074, using the splatEstimate() function. Specifically, the parameter “de.scale” was set to 1.0 and 1.2 to generate datasets with low and high clustering signals, respectively. Additionally, the parameter “batch.facScale” was set to 0.01 and 0.1 to generate datasets with low and high levels of differences between batches. Each simulated dataset consisted of 2000 genes and 1000 cells. The total number of groups in all simulation experiments was fixed at 8, and the number of cells in each group was randomly determined. To ensure variability, we generated 10 pairs of datasets for each experiment setting by altering the random seed.

### Differential expression analysis

Datasets are first integrated using Seurat V3 with top 4000 features, and then the clustering results of scDILT is added to the Seurat object as meta data. Cell type markers are obtained in the target cell cluster by comparing to the rest of clusters using Wilcoxon test. To understand these markers in a functional enrichment, we rank them by the log fold change from the DE analysis and used it for the gene set enrichment analysis (GSEA) implemented by the package fgsea[28] (1.19.4) in R.

## RESULTS

### Simulation

The performance evaluation of data integration methods involves assessing the ability of the integrated data to cluster cells of the same cell type across different datasets. To measure the clustering accuracy, we utilize two metrics: Adjusted Rand Index (ARI) and Normalized Mutual Information (NMI). Higher scores indicate better clustering accuracy. In order to assess the performance of scDILT under various scenarios, we simulate four datasets with two levels of clustering signals (CS) and two levels of batch effects.

To compare the clustering performance of scDILT with other state-of-the-art methods, including Seurat V3, CarDEC, and DESC, we conduct a comparative analysis. For CarDEC and DESC, we use the default parameters except for tuning the resolution parameter in DESC to achieve a similar number of clusters as the true labels. For Seurat V3, we employ the anchor-based integration method to integrate two batches of data, followed by clustering analysis on the integrated data.

The simulation results are summarized in Fig. 2. It is evident that scDILT outperforms all competing methods with the highest ARI and NMI, particularly in scenarios with low clustering signals (CS = 1.0). Seurat V3 achieves the second-best NMI and ARI scores across all simulated datasets, followed by Cardev and DESC. These results demonstrate that scDILT exhibits robust performance that is unaffected by low clustering signals and high batch effects.

**Fig 2.**
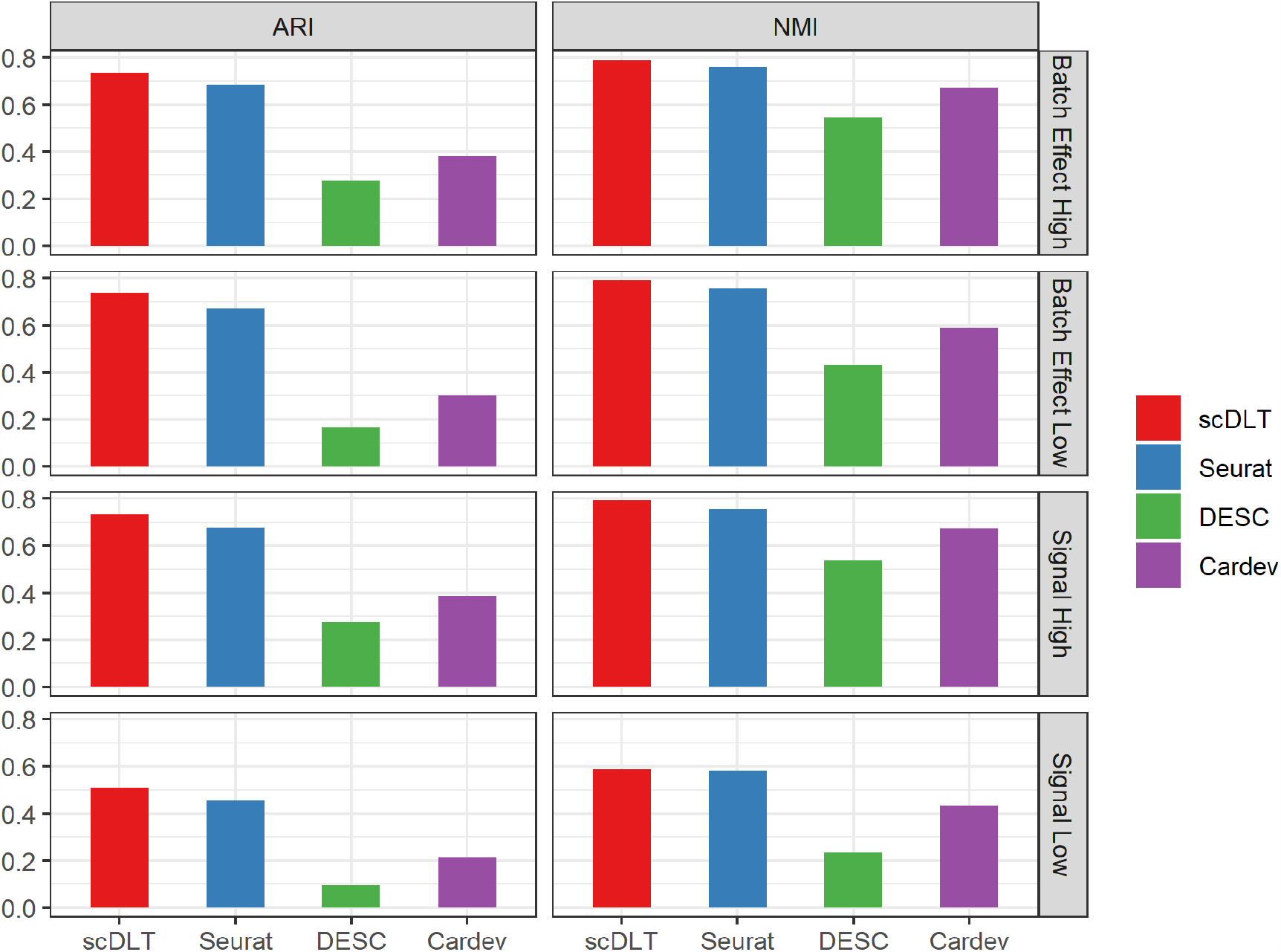
Clustering performance of scDILT and the competing methods in the simulation data with different settings. 1) high and low batch effect; 2) high and low clustering signal.

#### Integration of data in the same sample but different batches

In the first real data experiment, we focus on integrating two batches of scRNA-seq data obtained from the spleen of two mice, SLN111 and SLN208. Our assumption is that the cell type composition and proportion are nearly identical between the two batches of data. For this experiment, we designate the first batch of data as the reference dataset, which contains the true labels for the cell types. We utilize these reference labels to guide the clustering process for all cells, including those from both batches. By leveraging the known labels from the reference dataset, we aim to ensure accurate and consistent clustering across both batches of data.

In Fig. 3a, the t-SNE plots display the latent representations generated by scDILT, CarDEC, DESC, and Seurat V3 (from top to bottom) for the SLN111 dataset. The corresponding clustering performance of each method is depicted in Fig. 3b. Upon analysis, it is observed that scDILT achieves the highest ARI score (0.618) and the second-highest NMI score (0.625) among the competing methods, while the highest NMI score is 0.631. These results indicate that scDILT outperforms the other methods in terms of clustering accuracy and demonstrates its efficacy in integrating and analyzing the SLN111 dataset. In the t-SNE plot generated by DESC, cells from different batches appear to be completely separated. This observation suggests that the differences between the two batches overshadow the variations among different cell types. Conversely, in the t-SNE plots of Seurat V3, CarDEC, and scDILT, cells from different batches are mixed together. Each cell type contains cells from both batches, indicating that the variations among cell types play a more prominent role in the clustering process. However, it is worth noting that while CarDEC and Seurat V3 can effectively remove the batch effect between the two datasets, they struggle to separate different cell types in the integrated latent representation (as shown in Fig. 3a). Multiple clusters within different cell types appear to be connected to each other, making it challenging to discern distinct cell types using these clustering algorithms. In contrast, scDILT demonstrates superior performance by simultaneously separating different cell types in the latent space while effectively removing the batch effect. This results demonstrate the enhanced clustering accuracy of scDILT compared to the competing methods.

**Figure 3.**
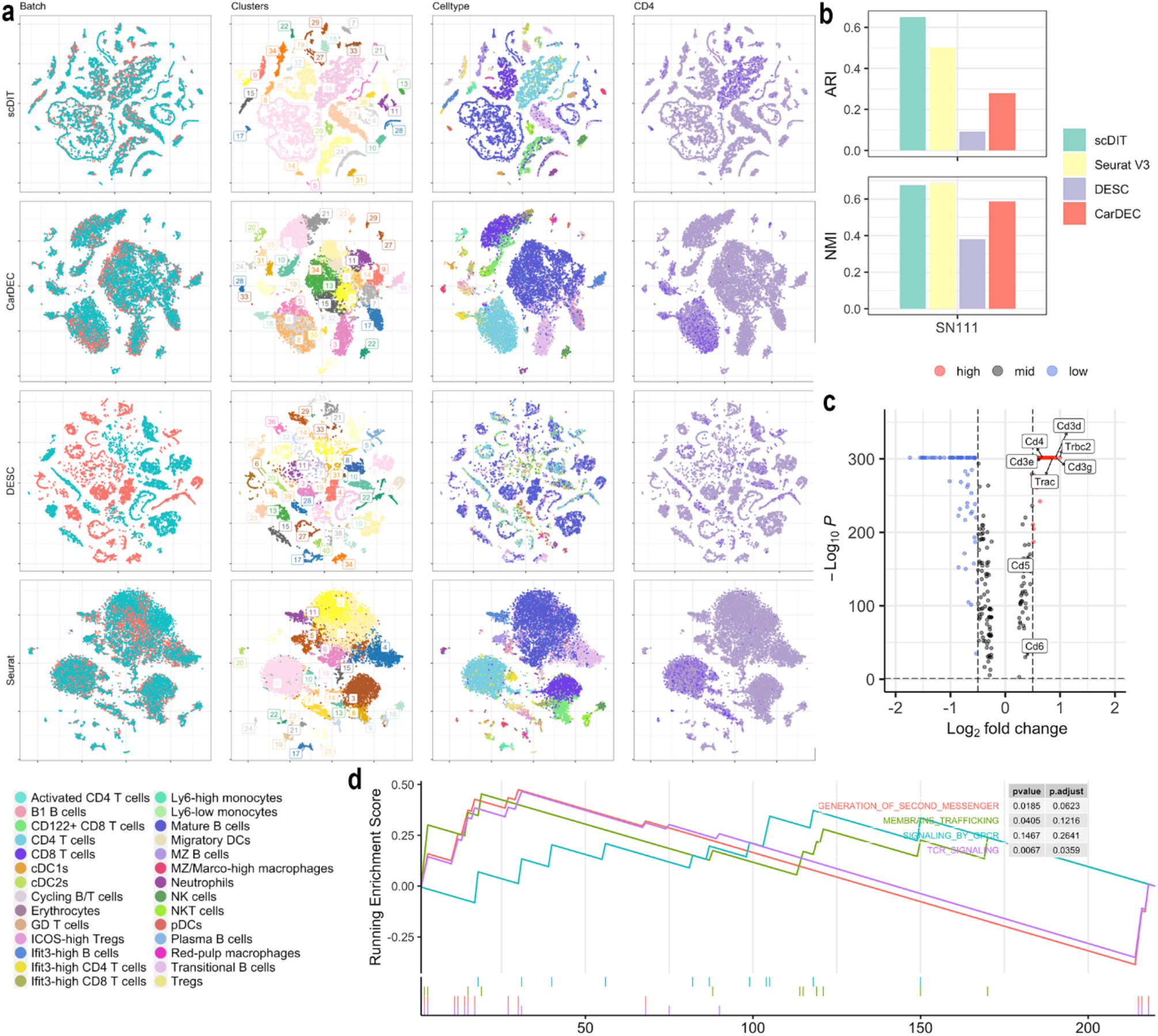
a. Latent representation of SLN111 dataset from scDILT and the competing methods. From top to bottom, t-SNE plots are listed from the latent representation from scDILT, CarDEC, DESC and Seurat V3 respectively. From left to right, cells are colored by the batches, clusters, celltypes and gene CD14 expression. b. Clustering performance of all methods with ARI on the top and NMI at the bottom. c. Volcano plot from the DE between cluster 14 (CD4 T cells) and the other clusters. d. GSEA plot of top 5 enriched pathways in geneset C8 with p value and p-adjusted value listed on the right side.

To further validate the clustering results obtained by scDILT, we employ differential expression (DE) analysis and gene set enrichment analysis (GSEA). Specifically, we perform DE analysis on cluster 14 by comparing its gene expression to that of all the other clusters using the Wilcoxon test. Given the clear observation in Fig. 3a that most cells in cluster 14 correspond to CD4 T cells, we can confidently assign cells of the same cell type from different batches into the same cluster post-integration. Consequently, we select CD4 as a marker gene and generate a plot illustrating its expression pattern within the t-SNE plots. This visualization helps to reinforce the accurate clustering and integration of CD4 T cells achieved by scDILT. In the t-SNE plots of scDILT, CarDEC, and Seurat V3, CD4 is mainly expressed in the cluster of CD4 T cells. However, in the t-SNE plot of DESC, CD4 expressed in multiple clusters. Besides CD4, multiple T cell marker genes, including CD27[29], CD28[30], LCK[31], TCF7[32], CD3D[33] show high log2 fold change (>2) and small P-value (<0.05) (Fig. 3c) in the result of scDILT. We then perform a GSEA by using the DEGs in the CD4 T cells. The gene set CP REACTOME from MSigDB Collections (cell type signatures) is used. In the top 10 significant pathways (NES > 2, pval < 0.05), 4 of them are T Cells related pathways^37^, including “Generation of second messenger”, “Membrane trafficking”, “Signaling by GPCR”, and “TCR signaling” (Fig. 3d). The “Generation of second messenger” pathway involves the production of molecules that transmit signals within the cell^38^(Table S1). “Membrane trafficking” enables the movement of proteins and other molecules between cellular compartments, crucial for T cell activation and antigen presentation^39^. “Signaling by GPCR” refers to the activation of G protein-coupled receptors, which initiate intracellular signaling cascades regulating T cell migration and activation^40^. “TCR signaling” pathway plays a central role in T cell activation by transducing signals from the T cell receptor, leading to diverse cellular responses^38^. These pathways collectively contribute to the intricate processes involved in T cell biology and immune responses. These results consolidate the clustering results of scDILT.

Fig. 4 illustrates the results obtained for dataset SLN208. In this dataset, scDILT demonstrates superior performance compared to the competing methods in terms of removing batch effects, separating cell types, and overall clustering performance (Fig. 4a-b). Cluster 11 is selected for further downstream analyses based on the obtained results. The cells in this clusters are mature B cells. Here we illustrate the expression of EBF1[34] which is a marker gene of multiple cell types including the B doublets, transitional B cells, lfit3-high B cells and so on. EBF1 expressed in multiple clusters in the t-SNE plots of scDILT and DESC but concentrated in a large cluster in the t-SNE plots of CarDEC and Seurat V3. This result indicates that scDILT and DESC can separate the B cell sub-types after integration in the latent representation. Other marker genes BCL11A, CD79B, CD74, CD19, CD55[34] and MS4A1 are also highly expressed in the cluster 11 from the results of scDILT (Fig. 4c). Besides, in the results of GSEA of the cluster 11, 4 out of 10 top enriched pathways are B cell type related^37^, including “MHC class II antigen presentation”, “Antigen activates B cell receptor BCR leading to generation of second messengers”, “Signaling by the B cell receptor BCR”, and “Interferon gamma signaling”(Fig. 4d, Table S2). The “MHC class II antigen presentation” pathway describes the presentation of antigens by major histocompatibility complex class II molecules, essential for B cell activation and antigen recognition^41^. The “Antigen activates B cell receptor BCR leading to generation of second messengers “pathway highlights the activation of B cell receptors (BCRs) by antigens, leading to the generation of second messengers such as cAMP, which regulate various cellular processes^42^. The “Signaling by the B cell receptor BCR “pathway details the intracellular signaling cascades initiated upon BCR activation, triggering B cell proliferation, differentiation, and antibody production^43^. The “Interferon gamma signaling “pathway involves the signaling of interferon-gamma, a cytokine that influences B cell activation, antibody class switching, and immune responses^44^. These pathways collectively contribute to the orchestration of B cell-related immune processes. To sum up, the results in this part demonstrate that scDILT has a superior performance in data integration and label transferring for the datasets in the same tissue but sequenced in different batches.

**Figure 4.**
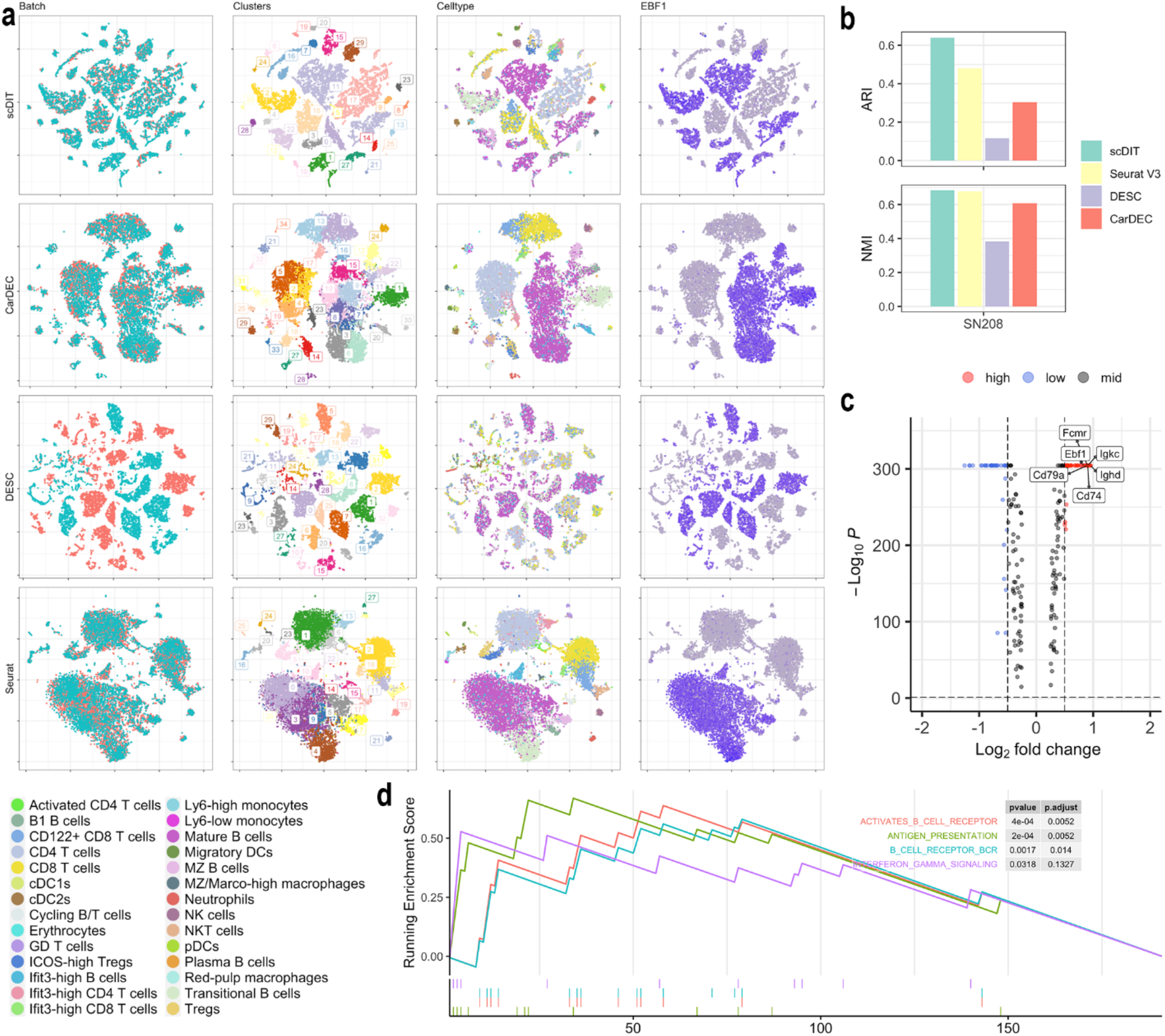
a. Latent representation of SLN208 dataset from scDILT and the competing methods. From top to bottom, t-SNE plots are from scDILT, CarDEC, DESC and Seurat V3 respectively. From left to right, cells are colored by the batches, clusters, celltypes and gene CD79a expression. b. Clustering performance of all methods with ARI on the top and NMI at the bottom. c. Volcano plot from the DE between cluster 11 (mature B cells) and the other clusters. d. GSEA plot of top five enriched pathways in geneset C8 with p value and p-adjusted value listed on the right side.

#### Integration of data in the same tissue but different batches

In this experiment, we performed data integration on scRNA-seq data obtained from different batches of human lung samples. The first batch, denoted as GSE131907, consisted of 208,506 cells derived from 58 lung tissues of 44 patients, while the second batch, denoted as GSE123904, comprised 40,505 cells from 17 lung tissues of 15 patients. Due to the large size of these datasets, we randomly selected 500 cells from each cell type per batch for analysis. Although these cells originated from the same tissue, their cell type compositions were not identical. Out of the 32 identified cell types, 11 were shared between the two datasets, 15 were unique to the first dataset, and 6 were unique to the second dataset. The first dataset served as the reference, containing the ground truth labels. The results of this experiment are depicted in Fig. 5, where scDILT exhibited the highest ARI and NMI among all methods (Fig. 5b). Conversely, in the t-SNE plots of DESC, cells were segregated into numerous small clusters, with multiple cell types emerging in each cluster. In contrast, the t-SNE plot of scDILT, CarDEC, and Seurat V3 showed well-separated clusters of cells with varying sizes. Notably, scDILT successfully preserved the unique clusters for individual datasets, aligning with the ground truth. On the other hand, CarDEC and Seurat V3 failed to accurately separate certain cell types in the latent representation, resulting in t-SNE plots containing a large ‘cluster’ comprising several smaller clusters and multiple cell types. Only scDILT demonstrated the ability to effectively separate all clusters in the t-SNE plot while maintaining distinct clusters for each dataset. For example, cluster 35 is mesothelial cells unique in the second dataset; and cluster 25 is tS1 cells unique in the first dataset. We select cluster 5 (mast cell) for the downstream analyses. We show the expression of marker gene KIT[35] in Fig. 5a and find that scDILT cluster most cells from different batches into a single cluster. The other two marker genes of mast cells, GATA2 and MS4A2[35], are also highly expressed in cluster 5 as shown in the volcano plot (Fig. 5c). In the results of GSEA, 4 out of 10 enriched pathways are mast cells related, including “travaglini lung basophil mast 2 cell”, “travaglini lung basophil mast 1 cell”, “cui developing heart C7 mast cell”, and “durante adult olfactory neuroepithelium mast cells” (Fig. 5D, Table S3). These mast cells play important roles in various physiological processes, including immune responses, tissue homeostasis, and sensory perception. The specific pathways involved in these cell types may include processes related to cell activation, degranulation, immune signaling, cytokine production, and tissue-specific functions. These findings provide compelling evidence of the superior performance of scDILT in effectively integrating datasets derived from the same tissue but originating from different patients. The ability of scDILT to successfully merge and harmonize these datasets highlights its robustness and effectiveness in overcoming inter-patient variability and capturing the underlying biological heterogeneity. By accurately integrating and aligning the data, scDILT enables a comprehensive analysis of the shared and unique features across patients, thereby facilitating the identification of key biological insights and potential biomarkers associated with the specific tissue of interest.

**Figure 5.**
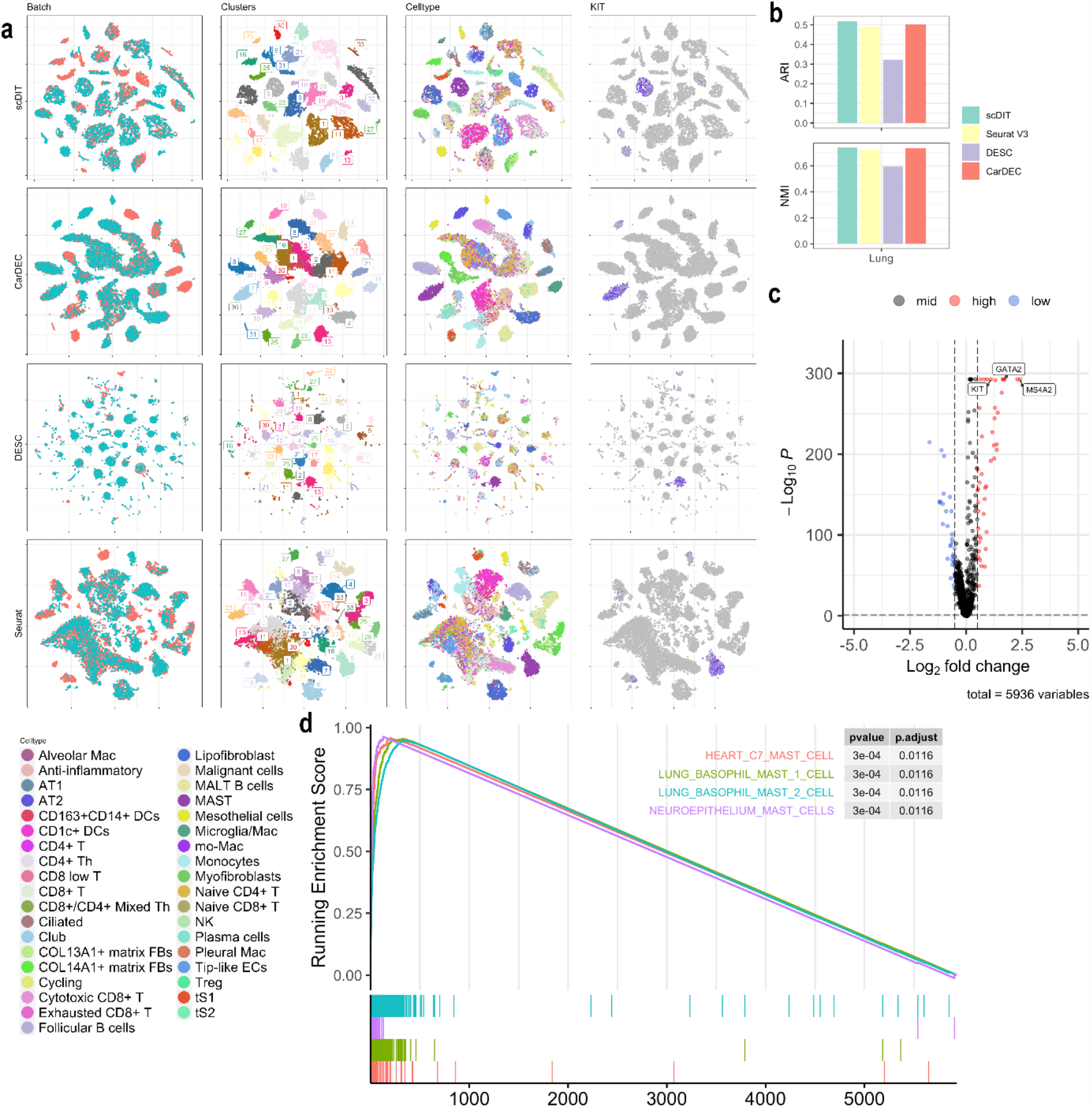
a. Latent representation of Lung cancer dataset from scDILT and the competing methods. From top to bottom, t-SNE plots are from scDILT, CarDEC, DESC and Seurat V3 respectively. From left to right, cells are colored by the batches, clusters, celltypes and gene KIT expression. b. Clustering performance of all methods with ARI on the top and NMI at the bottom. c. Volcano plot from the DE between cluster 5 (mast cells) and the other clusters. Genes are listed within the box. d. GSEA plot of top 5 enriched pathways in C8 geneset with p value and p-adjusted value listed on the right side.

#### Integration of data in the sample with different treatments

In the third experiment, we integrate the scRNA-seq of human PBMC with different treatments. We assume that there are some differences of cell type composition and/or proportion among different datasets. We use four datasets in this experiment including: 1) PBMC from a healthy human; 2) PBMC from a patient with Drug-induced hypersensitivity syndrome/drug reaction with eosinophilia and systemic symptoms (DiHS/DRESS); 3) PBMC from a DiHS/DRESS patient with sulfamethoxazole (SMX) and trimethoprim (TMP) treatment; 4) PBMC from a DiHS/DRESS patient with SMX, TMP, and tofacitinib (TOFA) treatment. Since only the healthy PBMC dataset has true labels, we this dataset as the reference to guide the clustering of the other datasets. Fig. 6 displays the t-SNE plots of the latent representations obtained from scDILT and Seurat V3 on the PBMC datasets. Seurat V3 effectively removes the batch effect, but it fails to clearly distinguish between various cell types in the latent space. In contrast, scDILT demonstrates a satisfactory performance in both batch effect removal and cell type separation. Notably, the well-separated cell types in the latent space of scDILT allow for the observation of variations in cell type composition across different batches and the identification of distinct batch proportions among different cell types. In Fig. 6, the cells from the healthy PBMC sample are colored based on their true labels/cell types, while the remaining cells are represented in blue. Interestingly, two new clusters, namely cluster 1 and 2, are identified in the patient samples, while absent in the healthy PBMC data. Notably, cluster 1 and 2 exclusively contain cells from patients, with cluster 2 lacking cells from the TOFA treatment group. Conversely, cluster 12 exclusively comprises cells without any treatments and represents the CD14+ monocyte cell type. These findings suggest that treatments such as SMX, TMP, and/or TOFA have the potential to eliminate CD14+ monocytes in the peripheral blood of patients. In summary, this experiment demonstrates that scDILT has a superior performance in batch effect removal and cell type separation, providing a chance to find new cell types in the samples with different interventions.

**Figure 6.**
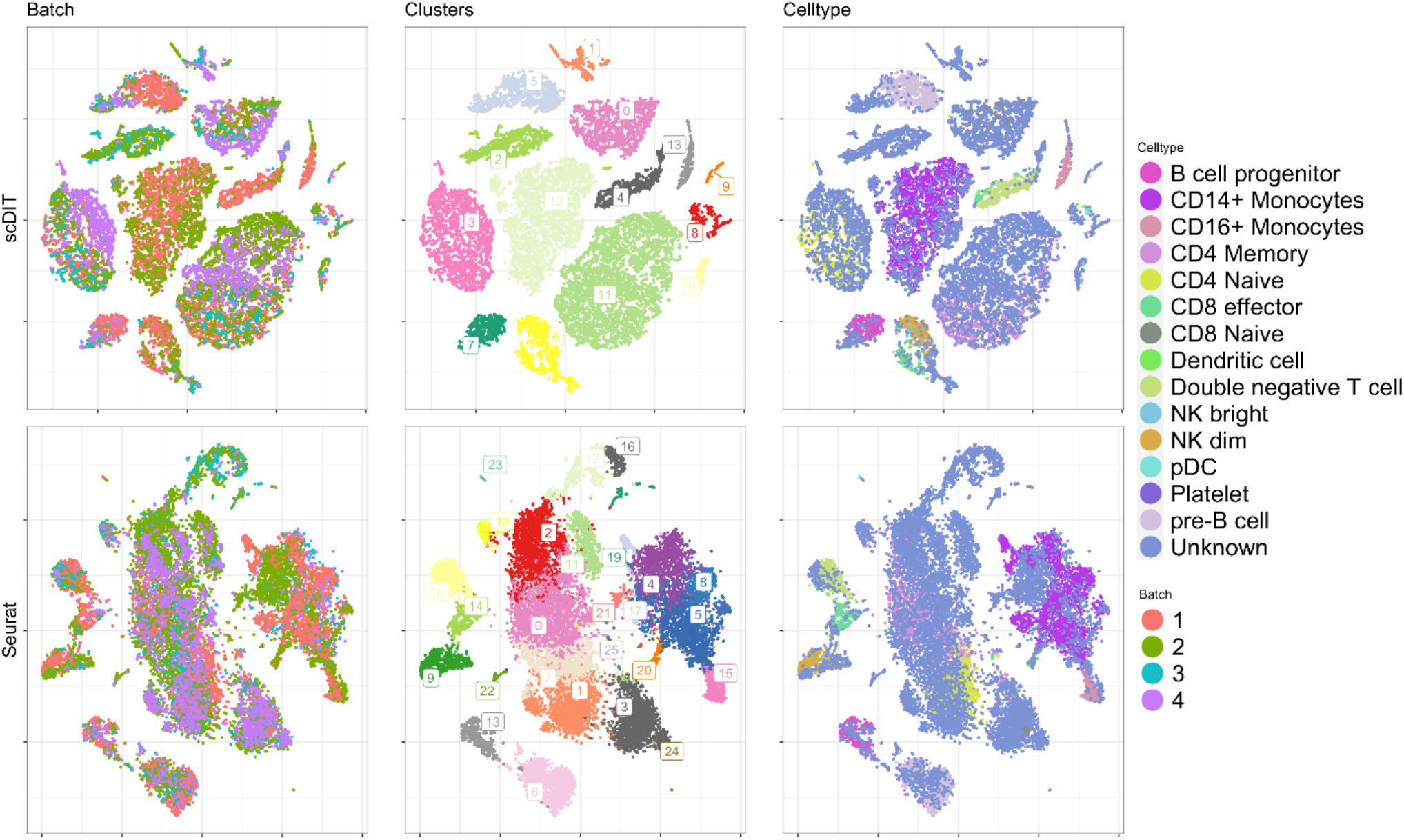
Latent representation of PBMC dataset with different treatments from scDILT and the competing methods. From top to bottom, t-SNE plots are from scDILT and Seurat V3 respectively. From left to right, cells are colored by the batches, clusters and celltypes. Legends of cell types and batch are attached to the right. Batch information: 1) PBMC from a healthy human; 2) PBMC from a patient with Drug-induced hypersensitivity syndrome/drug reaction with eosinophilia and systemic symptoms (DiHS/DRESS); 3) PBMC from a DiHS/DRESS patient with sulfamethoxazole (SMX) and trimethoprim (TMP) treatment; 4) PBMC from a DiHS/DRESS patient with SMX, TMP, and tofacitinib (TOFA) treatment.

### Integration of scRNA-seq and scATAC-seq data

In the final experiment, we integrate the Single-cell Multimode ATAC Gene Expression (SMAGE-seq) dataset of human PBMC obtained from the 10X Genomics website. To integrate the multimodal data, we perform separate integration for the mRNA and ATAC modalities. The results of mRNA integration are presented in Fig. 7, while the results of ATAC integration are shown in Fig. 8. Notably, scDILT outperforms the other methods with the highest ARI and NMI scores for both scRNA-seq and scATAC-seq data (Fig. 7b and Fig. 8b). Although all methods effectively remove the batch effect (Fig. 7a), certain challenges arise in the t-SNE plots. CarDEC fails to distinguish between CD4 memory cells and CD8 naive cells in the mRNA data, DESC separates CD14+ monocytes and CD4 memory cells into multiple clusters, and Seurat struggles to separate pre-B cells and B cell progenitors. However, scDILT successfully separates pre-B cells and B cell progenitors, albeit with slight overlap between CD4 memory cells and CD8 naive cells. In the t-SNE plot of the ATAC data, DESC fails to integrate the two batches, Seurat and CarDEC remove batch effects but struggle to separate different cell types such as B cell progenitors and platelet cells in the latent representation. Only scDILT accurately separates these cell types while effectively removing batch effects. Furthermore, we assess the expression of the CD14 marker gene in CD14 monocyte cells using t-SNE plots, and confirm its exclusive expression in the CD14 monocyte cluster. Other marker genes for CD14 monocytes, including SAT1, ZEB2, LYZ, CD36, and CD74^36^, also exhibit high expression in cluster 11, as identified by scDILT (Fig. 7C). In the pathway analysis, 3 out of top 10 enriched pathways are monocytes related, including “lung classical monocyte cell”, “lung nonclassical monocyte cell” and “lung ORL1 classical monocyte cell”(Fig. 7d, Table S4). The “lung classical monocyte cell” pathway refers to the molecular processes specific to classical monocytes in the lung, involving their recruitment, migration, antigen presentation, phagocytosis, cytokine production, and interactions with other immune cells. The “lung nonclassical monocyte cell” pathway represents the characteristics and functions of nonclassical monocytes involved in immune surveillance and tissue homeostasis. The “lung ORL1 classical monocyte cell” pathway describes the molecular features and activities of classical monocytes expressing the ORL1 receptor in the lung, which may include unique signaling events and functional roles in immune responses. In summary, this experiment demonstrates that scDILT has a superior performance in batch effect removal and cell type separation in not only scRNAseq data, but also scATAC-seq data, providing a chance to integrate multi-omics datasets.

**Figure 7.**
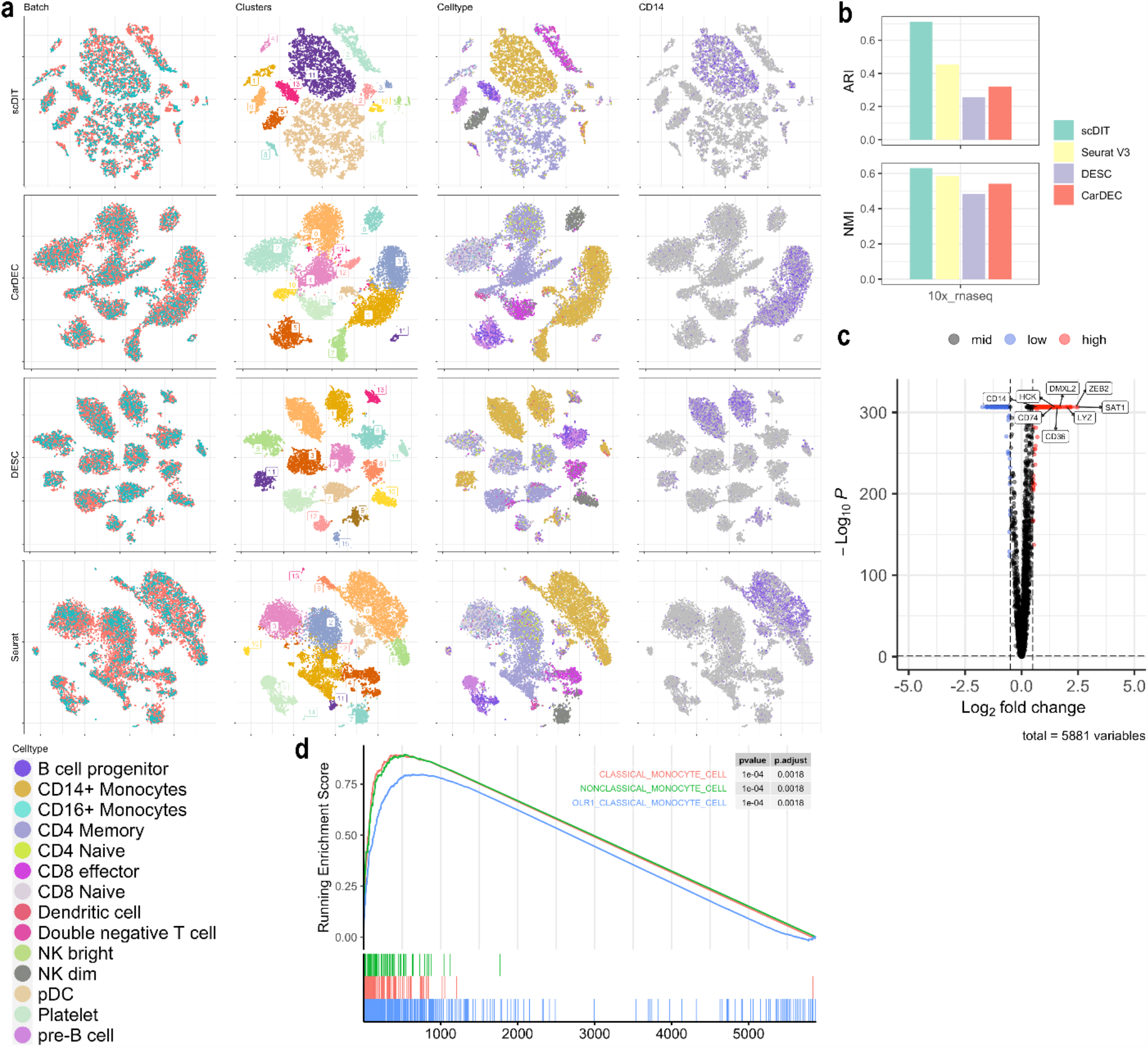
a. Latent representation of 10X granulocyte scRNAseq dataset from scDILT and the competing methods. From top to bottom, t-SNE plots are from scDILT, CarDEC, DESC and Seurat V3 respectively. From left to right, cells are colored by the batches, clusters, celltypes and gene CD14 expression. b. Clustering performance of all methods with ARI on the top and NMI at the bottom. c. Volcano plot from the DE between cluster 11 (CD4 monocytes) and the other clusters. d. GSEA plot of top 5 enriched pathways in C8 geneset with p value and p-adjusted value listed on the right side.

**Figure 8.**
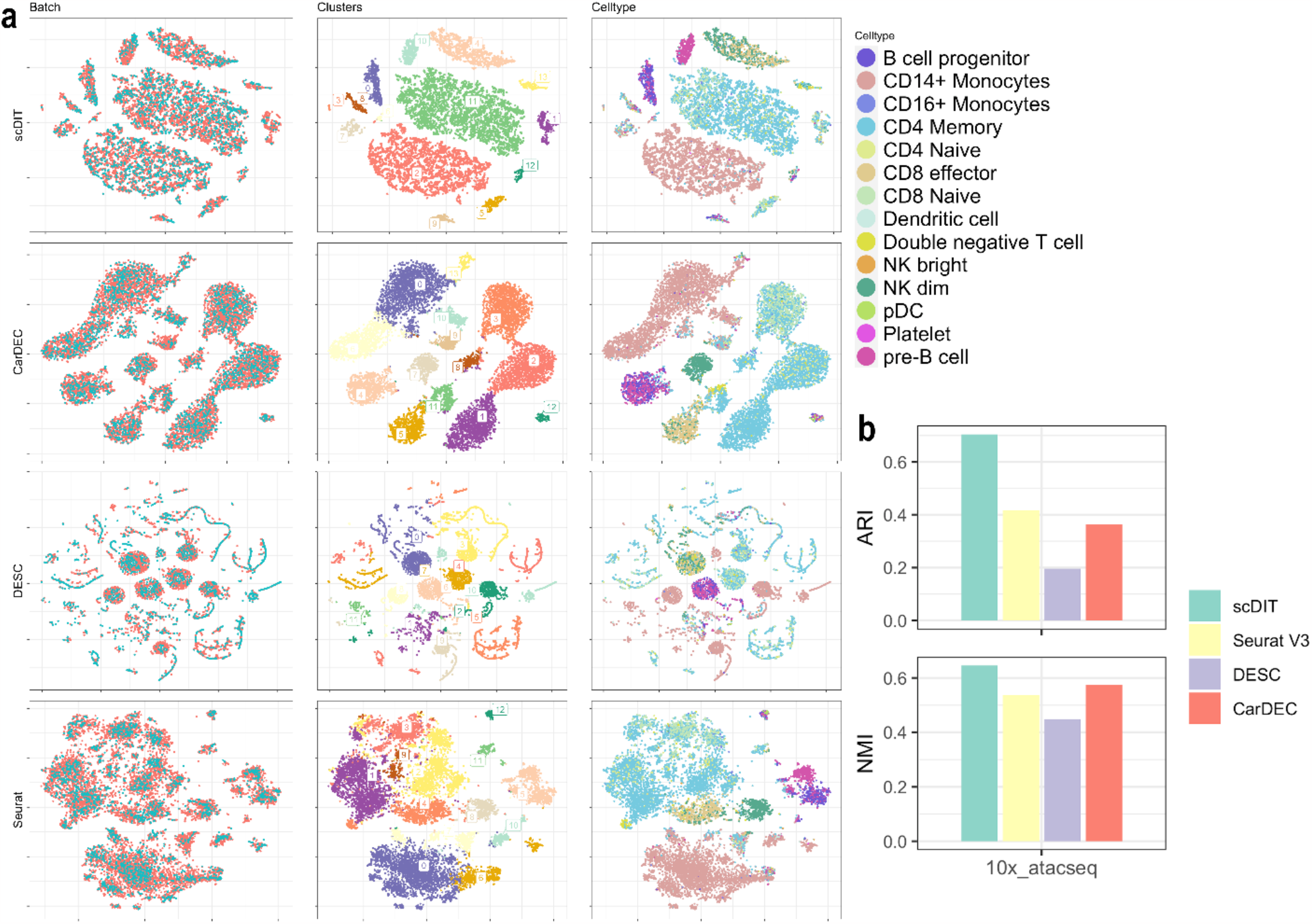
a. Latent representation of 10X granulocyte scATACseq dataset from scDILT and the competing methods. From top to bottom, t-SNE plots are from scDILT, CarDEC, DESC and Seurat V3 respectively. From left to right, cells are colored by the batches, clusters and celltypes. b. Clustering performance of all methods with ARI on the top and NMI at the bottom.

## Model test

### Time complexity

We performed simulations using various datasets with an increasing number of cells to investigate the relationship between cell numbers and the running times of scDILT. In Fig. 9a, we demonstrate that scDILT’s running time exhibits a linear increase with the number of cells. Specifically, for a given dataset X with N cells, we combined two instances of X to create a larger dataset with 2N cells, simulating the integration process. Notably, when integrating a large dataset with 2x100,000 cells, scDILT completes all tasks, including integration and clustering, in approximately 50 minutes. It is worth mentioning that scDILT is trained using batches of samples, with a batch size of 256, leading to a low space complexity even when working with large datasets.

**Figure 9.**
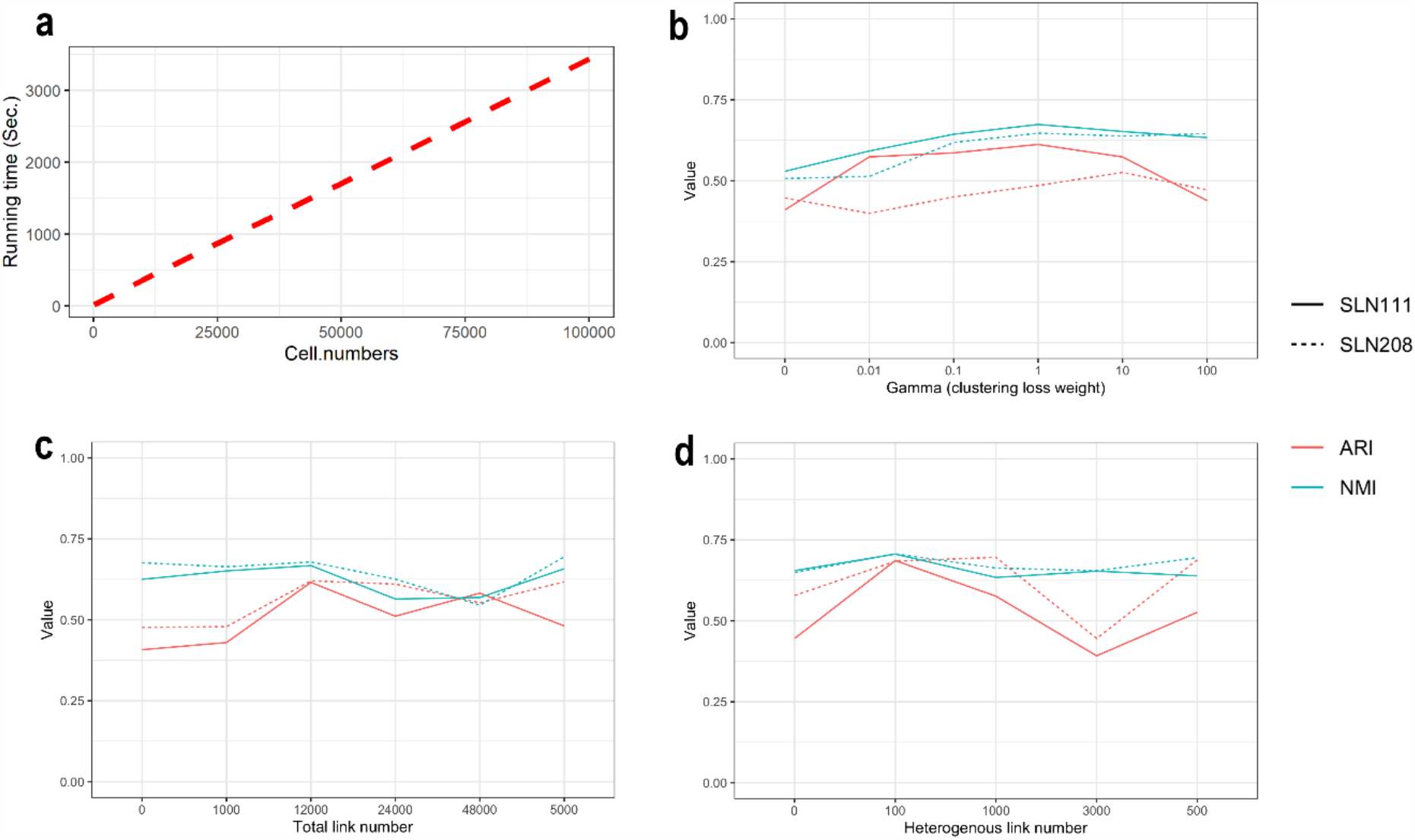
a. Running time of scDILT. We integrate two datasets with cell numbers 100, 500, 1000, 5000, 10000, 25000, 50000, 76500 and 100000. Running time of scDILT increases linearly with the increase of cell number. b. Relationship of γ (weight of clustering loss) and clustering performance. We set different γ value of 0, 0.01, 0.1, 1, 10, and 100. c. Relationship of total link number and clustering performance. We set different number of total links at 0, 1000, 5000, 12000, 24000, and 48000. d. Relationship of heterogenous link number and clustering performance. We set different number of heterogenous link at 0, 100, 500, 1000, and 3000. For panel b, c, and d, all clustering performance is evaluated using ARI (red) and NMI (blue) on two datasets SLN111 (solid line) and SLN208 (dashed line).

### Robustness tests

All model tests for scDILT were conducted using two spleen lymph node datasets (SLN111 and SLN208). The clustering loss weight, gamma, is an important parameter in our model. To evaluate the impact of the clustering loss on scDILT, we varied gamma values (0, 0.01, 0.1, 1, 10, and 100), while keeping other parameters fixed (Fig. 9b). We observed that when gamma is less than 1, the clustering loss enhances the clustering performance of scDILT. However, excessively high gamma values (>10) negatively affect scDILT’s performance. Additionally, we examined the contribution of homogeneous links to scDILT’s clustering performance. By increasing the total number of links (0, 1000, 5000, 12000, 24000, and 48000), while maintaining the ratio of 3-fold cross-links to ML links constant (Fig. 9c), we found that an appropriate number of homogeneous links (approximately equal to the number of cells) significantly improves clustering performance, while excessive links lead to inferior results. This may be due to excessively tight linking, which hinders the clustering process. Finally, we evaluated the impact of heterogeneous links on scDILT’s clustering performance. By increasing the number of heterogeneous links (0, 100, 500, 1000, 3000), while keeping the sum of heterogeneous and homogeneous ML links constant (3000), with the number of homogeneous CL links fixed at 9000 (Fig. 9d), we observed that an appropriate number of heterogeneous links (<1000) enhances the clustering performance, while too many heterogeneous links have a negative effect.

## CONCLUSION

This article presents scDILT, a novel tool for integrating scRNA-seq data from different batches or experiments. scDILT utilizes a conditional autoencoder and deep embedding clustering to effectively remove batch effects and ensure accurate clustering patterns across datasets. The inclusion of intra- and inter-dataset constraints enables scDILT to preserve clustering patterns in the reference datasets and accurately map cells in query datasets to pre-defined and annotated clusters. The extensive experiments demonstrate the superior performance of scDILT in data integration, making it a valuable tool for integrating single-cell datasets across different conditions, interventions, and time points. Additionally, scDILT shows potential for integrating multi-omics data, opening new possibilities for comprehensive analyses in the era of expanding single-cell sequencing technologies and available datasets.

## DATA AND CODE AVAILABILITY

The codes of scDILT are available on GitHub: https://github.com/xianglin226/scDILT. The source data and scripts of scDIT are available on GitHub: https://github.com/Jianlan0816/scDILT.

## Notes

### Competing Interest Statement

The authors have declared no competing interest.

